# Oxidized RNA Bodies compartmentalize translation quality control in *Saccharomyces cerevisiae*

**DOI:** 10.1101/2020.08.05.232983

**Authors:** James S. Dhaliwal, Cristina Panozzo, Lionel Benard, William Zerges

## Abstract

Cytoplasmic RNA granules compartmentalize phases of the translation cycle. We previously reported on the localization of oxidized RNA in human cells to cytoplasmic foci called oxidized RNA bodies (ORBs). Oxidized mRNAs are substrates of translation quality control, wherein defective mRNAs and nascent polypeptides are released from stalled ribosomes and targeted for degradation. Therefore, we asked whether ORBs compartmentalize translation quality control. Here, we identify ORBs in *Saccharomyces cerevisiae* and characterize them using fluorescence microscopy and proteomics. ORBs are RNA granules that are distinct from processing bodies and stress granules. Several lines of evidence support a role of ORBs in the compartmentalization of central steps in the translation quality control pathways No-Go mRNA decay and ribosome quality control. Active translation is required by both translation quality control and ORBs. ORBs contain two substrates of translation quality control: oxidized RNA and a stalled mRNA-ribosome-nascent chain complex. Translation quality control factors localize to ORBs. Translation quality control mutants have altered ORB number per cell, size, or both. Therefore, ORBs are an intracellular hub of translational quality control.

## INTRODUCTION

Ribosomes can stall at oxidized bases and other lesions in a defective mRNA. This is problematic because it can generate truncated and aggregation-prone polypeptides and, thereby, impair cell physiology (1, 2). Consequently, defective mRNAs are recognized and cleared by the translation quality control (TQC) pathways, No-Go decay (NGD) and ribosome-associated quality control (RQC). Colliding ribosomes at a translation blockage on a defective mRNA are recognized whereupon NGD separates their subunits and targets the aberrant mRNA for degradation (3–7). RQC then extracts the nascent polypeptide from the 60S ribosomal subunit, and targets it for degradation by the ubiquitin-proteasome system (8–12).

DNA and protein quality control are localized to specialized intracellular compartments (13–15), and NGD has been proposed to occur in the cytoplasmic RNA granules, processing bodies (P-bodies) and stress granules (16, 17). However, the reported proteomes of P-bodies and stress granules do not contain translation quality control proteins, with the exception of Rqc2 in yeast stress granules (18, 19). Therefore, additional work is required to determine whether TQC is compartmentalized and, if so, the nature of the compartment(s) involved.

An avenue to explore RNA quality control was suggested by our identification of cytoplasmic bodies that contain oxidized RNA in human cells (20). These “oxidized RNA-containing bodies” (ORBs) were identified by immunofluorescence (IF) staining of HeLa cells using an antibody against an oxidized form of guanine, 8-oxoguanosine (8-oxoG). ORBs were shown to be distinct from P-bodies and stress granules and their composition and functions remain unknown.

Here, we identify ORBs in *S. cerevisiae*. Using IF-microscopy and proteomics, we provide the following four lines of evidence that ORBs compartmentalize NGD and RQC. Two substrates of NGD and RQC localize to ORBs: oxidized RNA and a stalled mRNA-ribosome-nascent chain complex (RNC). NGD and RQC proteins are localized to ORBs. Mutants deficient for NGD, RQC, or both have altered ORB numbers per cell. Active translation is required by these pathways and ORBs. Finally, results of proteomics reveal relationships between ORBs and both P-bodies and stress granules.

## RESULTS

### ORBs are Cytoplasmic RNA Granules in Yeast

To determine whether ORBs exist in *S. cerevisiae*, we visualized the *in situ* distribution of 8-oxoG by IF-microscopy. Cells from cultures in exponential growth phase showed the 8-oxoG IF signal in cytosolic foci that numbered 5.4 per cell and were 450 nm in diameter (Fig. 1A and B). While 8-oxoG can be in RNA, DNA and nucleotides (21), these foci contain oxidized RNA because they were eliminated by RNase A treatment of fixed and permeabilized cells, they stained positive for RNA with an RNA-specific fluorescent dye and they did not stain for DNA with DAPI (Fig. 1C-E).

**Figure 1.**
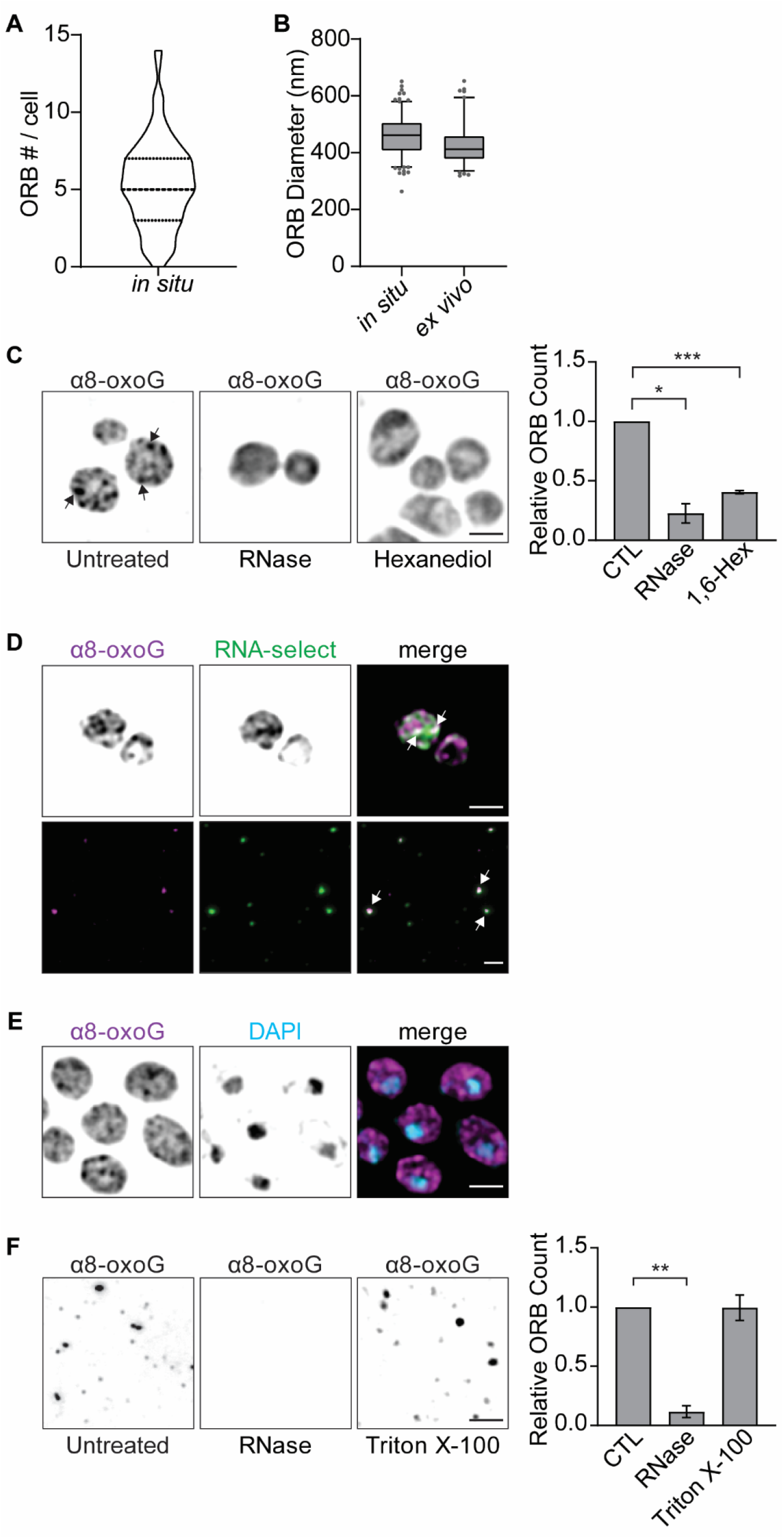
ORBs are an RNA granule in *S. cerevisiae*. (A) Violin plot of the ORB number per cell in unstressed cells from exponentially growing cultures. The median and interquartile ranges are indicated by thick and thin dashed lines, respectively (n=6). (B) Box plots of ORB diameter (nm) *in situ* and *ex vivo*. Box heights indicate the interquartile range, and whiskers indicate the 5^th^ and 95^th^ percentiles. (C) IF staining of exponentially growing unstressed cells revealed foci of 8-oxoG (black arrows). Cells were either untreated (CTL abbreviation) or treated with either RNase A, following fixation, or 1,6-hexanediol *in vivo*. Bar heights indicate ORB number per cell (relative to untreated cells) (n=3). Error bars = ± 1 SEM (D) Co-IF staining ORBs (α8-oxoG) and RNA with SYTO RNASelect *in situ* (top) and *ex vivo* (bottom). Arrows indicate foci of co-localized signals (E) Co-IF staining for ORBs (α8-oxoG) and DAPI. Scale bars = 2.0 μm. N values refer to the number of independent biological replicate experiments. (F) IF staining of ORBs (α8-oxoG) *ex vivo*. Lysates were untreated or treated with either RNase A or Triton X-100. Bar heights indicate ORB count (relative to untreated lysates) (n=3). Error bars = ± 1 SEM. (C and F) Statistical significance was determined by one-sample t-tests. * = p ≤ 0.05, ** = p ≤ 0.01, *** = p ≤ 0.001.

We asked whether ORBs are any of the membranous organelles that contain RNA and appear as foci in fluorescence microscopy. Cells were co-IF stained for 8-oxoG to visualize ORBs, and a GFP-tagged marker protein for mitochondria, Golgi, peroxisomes, endosomes, or autophagosomes (Fig. S1). In each case, ORBs tested negative revealing that they are distinct from these organelles and endosomes.

We then asked whether ORBs are RNA granules, i.e. membrane-less organelles that contain RNA and form by liquid-liquid phase separation (22). We observed that ORB number per cell was reduced by 40% during treatment *in vivo* with 1,6-hexanediol, which disrupts intermolecular interactions of liquid-liquid phase-separation that underlie RNA granule formation (Fig. 1C) (23, 24). Like P-bodies and stress granules, ORBs were stable in lysates, i.e. *ex vivo* (19,25) (Fig. 1F). These *ex vivo* foci were ORBs because they were similar in diameter, stained positive for RNA, and were eliminated by RNase A (Fig. 1B, D, and F). Finally, ORBs were resistant *ex vivo* to the non-ionic detergent Triton X-100, a property of RNA granules but not membranous organelles (26, 27) (Fig. 1F). Together, these results reveal that ORBs are RNA granules.

### ORBs are Affected by Translation Inhibition

Translational roles of P-bodies and stress granules were revealed, in part, by the effects of the translation inhibitors cycloheximide and puromycin (28, 29). Here, we found that treatment of cells with either drug reduced the number of ORBs per cell by over 2-fold (Fig. 2A and B). These results suggest a connection between ORBs and translation. However, ORBs are neither P-bodies nor stress granules because they lack canonical proteins of these RNA granules (Fig. S1).

**Figure 2.**
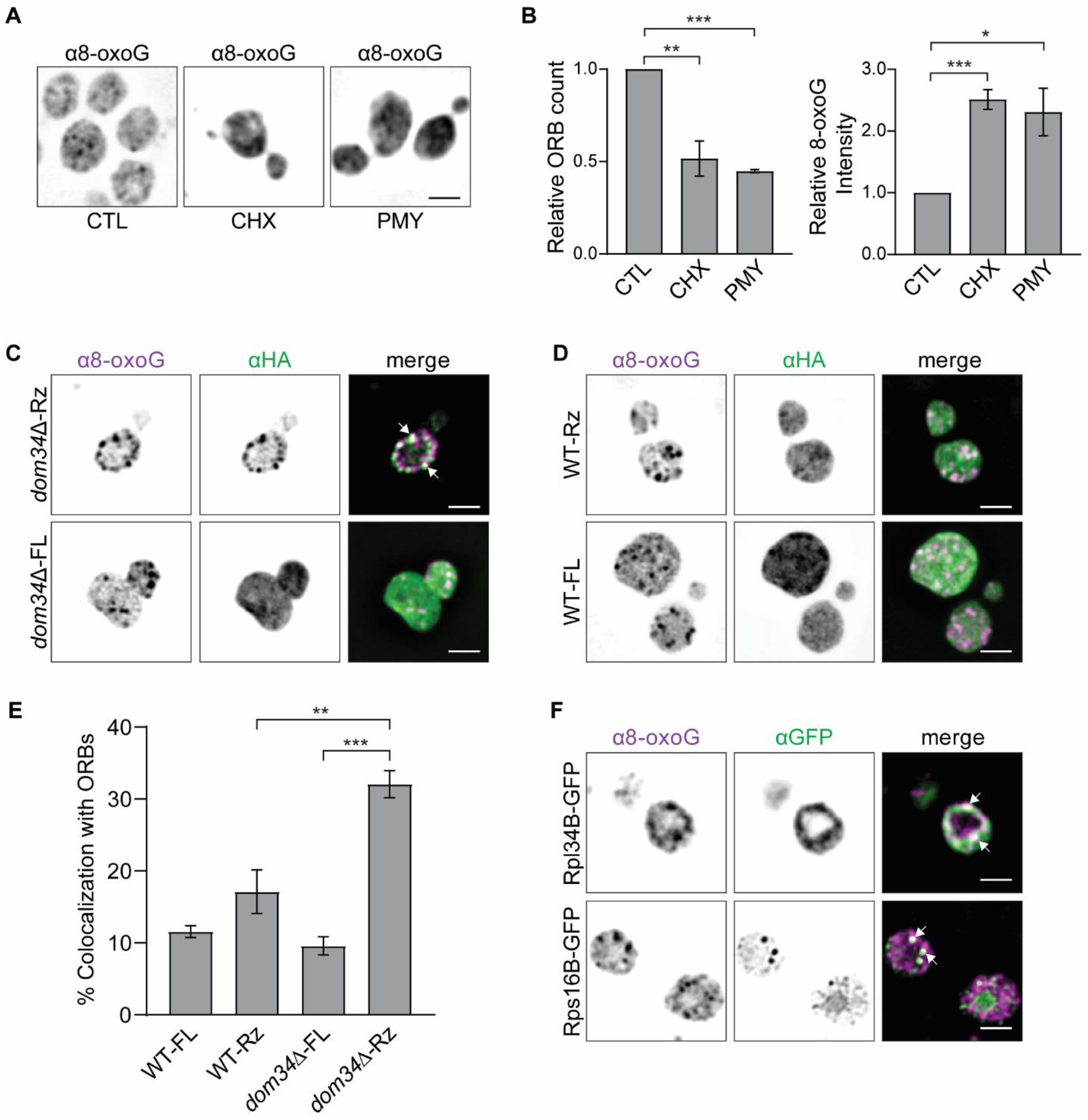
Stalled ribosome-nascent chain complexes localize to ORBs. (A) IF staining of ORBs (α8-oxoG) in cells that were untreated (CTL) or treated *in vivo* with either cycloheximide (CHX) or puromycin (PMY). (B) Bar heights indicate relative ORB number per cell (left) or average 8-oxoG intensity per cell (right) following treatment with CHX (n=5) or PMY (n=4). Error bars = ± 1 SEM. Statistical significance was determined by one-sample t-tests. (C and D) Co-IF staining of ORBs (α8-oxoG) and HA-Ura3, either as nascent polypeptides in the arrested RNC (Rz), or as full length and post-translation (FL). Genetic backgrounds were (C) *dom34*Δ, to stabilize the arrested RNC, or (D) WT, as the control. (E) Bar heights indicate the percentages of ORBs that colocalize with foci of HA-Ura3 (n =4 for WT-FL and WT-Rz, 5 for *dom34*Δ-FL, and 6 for *dom34*Δ-Rz). Error bars = ± 1 SEM. Statistical significance was determined by unpaired two-sample t-tests. (F) Co-IF staining of 8-oxoG with Rpl34B-GFP (top) or Rps16B-GFP (bottom). Arrows in C, D, and F indicate co-localized foci. Scale bars = 2.0 μm. N values refer to the number of independent biological replicate experiments. * = p ≤ 0.05, ** = p ≤ 0.01, *** = p ≤ 0.001.

We also observed that the intensity of the 8-oxoG IF signal throughout cells increased by over 2-fold during translation inhibition (Fig. 2B). This result is consistent with the known role of translation in clearing oxidized mRNAs through TQC pathways (2). Together, these results raised the possibility that ORBs compartmentalize TQC.

### A stalled Ribosome-Nascent Chain Complex Localizes to ORBs

If ORBs compartmentalize TQC, they should be enriched in a translationally arrested RNC, a TQC-specific substrate (30, 31). An arrested RNC was generated on a *URA3* mRNA with a self-cleaving hammerhead ribozyme inserted into its coding region (mRNA1Rz in Navickas et al., 2020). Self-cleavage generates a 3’ truncation which arrests translating ribosomes. Our rationale was to detect this arrested RNC *in situ*, relative to ORBs, by IF-staining an HA-epitope tag at the N-terminus of the nascent URA3 polypeptide. This was hampered by rapid disassembly of the arrested RNC by TQC. Indeed, the HA-Ura3 nascent polypeptide was not detected in a wild-type strain for NGD (Fig. S2). However, it was detected in *dom34*Δ background (Fig. S2), due to NGD deficiency (3). Moreover, it was shown previously that this truncated *URA3* mRNA in *dom34*Δ background is bound by an array of colliding ribosomes (31, 32). Therefore, the HA-Ura3 nascent polypeptide in *dom34*Δ background can serve as a marker for the arrested RNC *in situ.* Importantly, the use of a truncated mRNA prevents translation read-through and the generation of full-length HA-tagged polypeptide and background IF-signal, as occurs with many reporter constructs encoding stalling-prone defective mRNAs (33).

IF-microscopy images showed the HA signal from the arrested RNC localized to ORBs in *dom34*Δ background (Fig. 2C). In contrast, there was significantly less co-localization of the arrested RNC in the wild-type strain or of the full-length HA-Ura3 polypeptide in either background (Fig. 2C, D, and E). Therefore, this TQC-specific substrate localizes to ORBs.

Since ribosomes are present in arrested-RNCs, we asked whether ORBs contain ribosomal subunits by IF-staining GFP fused to ribosomal proteins. ORBs IF-stained for both ribosomal proteins tested; one of each of the ribosomal subunits (Fig. 2F). Additional ribosomal proteins are shown to localize to ORBs below. Therefore, ORBs contain both ribosome subunits, as would be expected for a location of TQC.

### TQC Factors Localize to ORBs

If ORBs compartmentalize TQC, they should have TQC proteins. To test this prediction, cells were IF-stained for the following GFP-tagged TQC factors: Hel2, which acts prior to NGD and RQC by recognizing colliding ribosomes, Hbs1 and Dom34, which act in NGD, and Rqc2, which acts in RQC (34, 35). Cells were co-IF stained for GFP and 8-oxoG to visualize GFP-tagged TQC proteins and ORBs, respectively. IF-microscopy images of these cells revealed that ORBs IF-stained for each of these TQC proteins (Fig. 3A). Quantification of the overlap of ORBs and foci with the GFP IF-signals was 24% for Dom34-GFP, 25% for Hbs1-GFP, 24% for Hel2-GFP, and 32% for Rqc2-GFP (Fig. 3, Table S1). ORBs also IF-stained for Dom34-GFP or Hbs1-GFP *ex vivo*, i.e. after the dispersal of intracellular material (Fig. 3B). Therefore, fortuitous overlap *in situ* is highly improbable. Our observation that only subpopulations of ORBs IF-stained for any particular protein was not unexpected because other translation-related RNA granules are similarly heterogeneous such that only subpopulations contain any particular protein or mRNA (36–39). Therefore, TQC proteins localize to ORBs.

**Figure 3.**
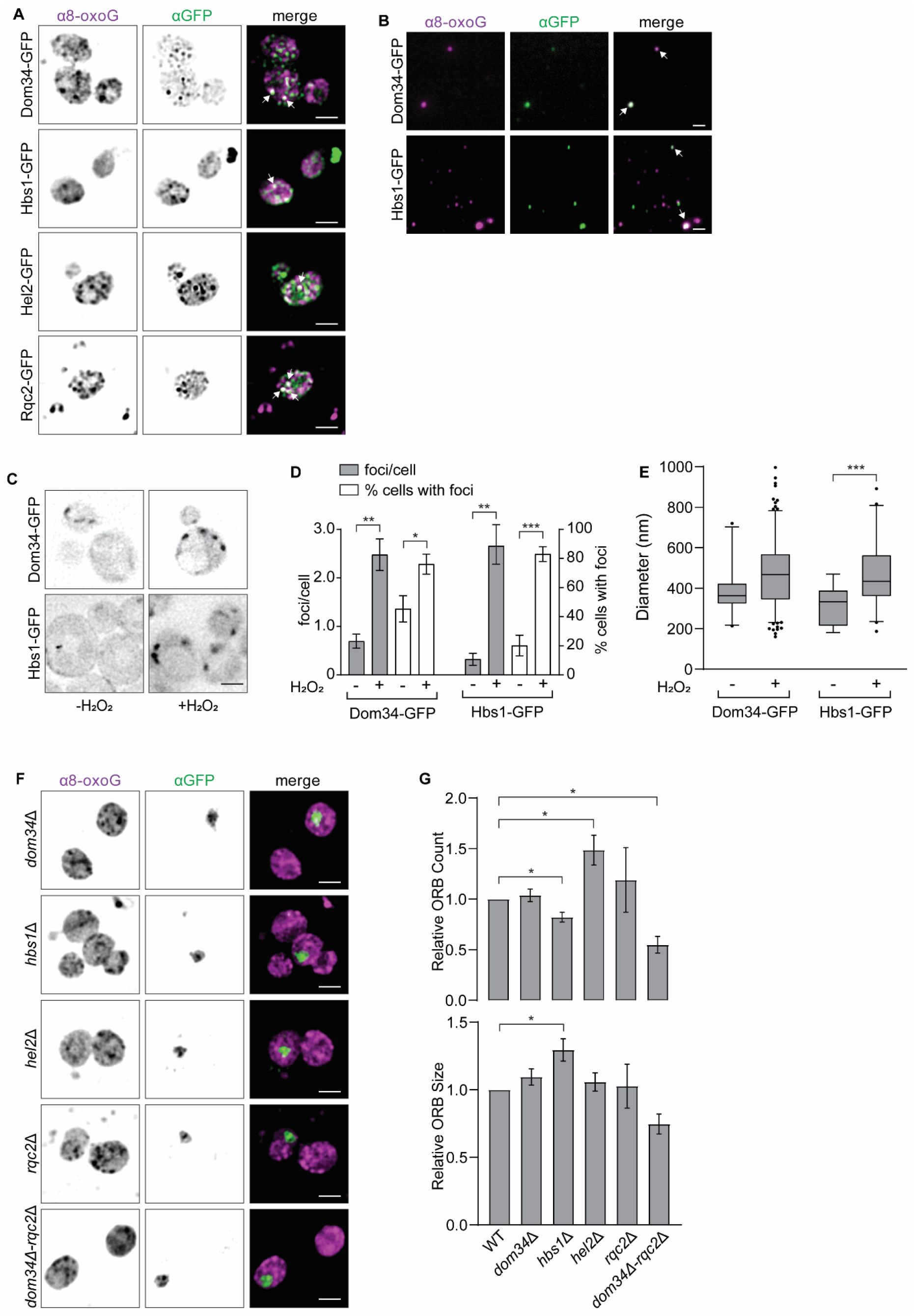
TQC factors localize to ORBs and are required for WT ORB number per cell. (A) Co-IF staining of ORBs (α8-oxoG) and the indicated GFP-tagged TQC proteins *in situ*. (B) Co-IF staining of ORBs (α8-oxoG) and the indicated GFP-tagged TQC proteins *ex vivo*. (C) Dom34-GFP and Hbs1-GFP fluorescence *in vivo* in cells without or with exposure to H_2_O_2_ (5.0 mM). (D) For Dom34-GFP or Hbs1-GFP *in vivo*, bar heights represent the number of foci per cell (left axis) and percentage of cells with foci (right axis), without or with exposure to H_2_O_2_ (n=4). Error bars = ± 1 SEM. Statistical significance was determined by unpaired two-sample t-tests. (E) Box plots of Dom34-GFP or Hbs1-GFP foci diameter (nm) without or with exposure to H_2_O_2_ (n=4). Box heights indicate the interquartile range, whiskers the 5^th^ and 95^th^ percentiles. Unpaired two-sample t-tests revealed no significant difference in diameter of Dom34-GFP foci without or with H_2_O_2_ treatment and a significant increase in diameter of Hbs1-GFP foci during H_2_O_2_ treatment. (F) Co-IF staining of ORBs (α8-oxoG) and GFP in mixed wild-type and mutant cells. Wild-type cells expressed Rpb2-GFP (a nuclear protein) which allowed them to be distinguished from mutant cells by IF-staining GFP. (G) Bar heights show (top) ORB number per cell and (bottom) ORB size relative to wild type in the TQC mutants for *dom34*Δ, *hbs1*Δ, and *hel2*Δ (n= 4). *rqc2*Δ and *dom34*Δ-*rqc2*Δ (n =3). Error bars = ± 1 SEM. Statistical significance was determined by unpaired two-sample t-tests. Arrows in A and B indicate co-localized foci. Scale bars = 2.0 μm. N values refer to the number of independent biological replicate experiments. * = p ≤ 0.05, ** = p ≤ 0.01, *** = p ≤ 0.001.

To determine whether, *in vivo,* Dom34 and Hbs1 localize to foci such as ORBs, we examined the distributions of their GFP-tagged versions in live cells by epifluorescence microscopy. Most cells from cultures in exponential growth phase showed Dom34-GFP or Hbs1-GFP throughout their cytoplasm, with only small minorities showing a focus (Fig. 3C and D). Under these non-stress conditions, only Dom34-GFP foci were more abundant and larger than foci of GFP alone, suggesting the natural occurrence of Dom34 foci (Fig. S3). This was not the case for Hbs1-GFP foci. When cells from the same cultures were exposed to the fixation conditions used for IF staining, we observed a significant increase in the number of foci of Dom34-GFP or Hbs1-GFP (Fig. S3). To determine whether the formation of these foci is a response to stress induced by formaldehyde, we examined cells from the same cultures following their exposure to H_2_O_2_, a ROS which causes oxidative stress (21). H_2_O_2_ treatment increased the number per cell of Dom34-GFP and Hbs1-GFP foci (Fig. 3C and D). This was surprising given that stress-induced localization of NGD factors to foci has not been reported previously.

The foci of Dom34-GFP or Hbs1-GFP that are induced by formaldehyde or H_2_O_2_ were likely the same structures because the effects of these stressors were not additive (Fig. S3). Furthermore, these foci are likely ORBs because they have the same diameter, in addition to having Dom34-GFP and Hbs1-GFP (Fig. 1B and 3E). Therefore, the formation of foci with translation quality control proteins in live cells provides validation of their detection in ORBs by IF-microscopy.

### Genetic Evidence for the Compartmentalization of TQC in ORBs

As another test of a role of ORBs in TQC compartmentalization, we asked whether their size and number per cell are altered in TQC-deficient mutants (Fig. 3F and G). Relative to wild type, *hel2*Δ cells had 50% more ORBs; *hbs1*Δ cells had 18% fewer ORBs, which were 30% larger; finally, *dom34*Δ*-rqc2*Δ double mutant cells had 46% fewer ORBs. The latter is a synthetic phenotype; it was not seen in *dom34*Δ or *rqc2*Δ single mutants. These phenotypes provide functional evidence that ORBs compartmentalize TQC.

### Proteomic Results Support ORBs as Being an RNA Granule that Compartmentalizes TQC

We used proteomics to explore ORB functions. ORBs were immunoaffinity-purified by a procedure developed for stress granules, but using the antibody against 8-oxoG (Fig. S4) (19). Proteomic analyses identified 822 candidate ORB proteins by at least two unique peptides, and which were not depleted by cycloheximide treatment (Fig 2 A and B, Table S1).

By co-IF staining 109 of these GFP-tagged candidate proteins with 8-oxoG *in situ*, we identified 60 proteins that localize to ORBs (Table S1) (Cdc42 was IF stained with an antibody against it and not a GFP-tagged version). These proteins were seen to localize to ORBs in microscopy images and tested positive for ORB-localization with a custom-written ImageJ macro (Table S1). Two proteins with high ORB-localization scores from the macro, Psa1-GFP and Nmd3-GFP, also localize to foci during *in vivo* treatment with formaldehyde, H_2_O_2_, or both (Fig. S3). To the list of validated ORB proteins we added 8 proteins that we seen to localize to ORBs prior to the proteomic analysis but, nonetheless, were supported by one unique peptide and non-depletion in the cycloheximide control. The resulting validated ORB proteome contains 68 proteins (Table S1).

This proteome supports ORBs as being RNA granules with functions in RNA metabolism. GO analysis revealed that, relative to the total yeast proteome, the ORB proteome is enriched in protein classes found in cytoplasmic RNA granules: e.g. ATPases, RNA-binding proteins, and RNA-helicases, (Fig. 4A) (18, 19). We also identified RNA-binding proteins by comparison to published databases that contain RBPs not found by GO analysis (Table S1) (40, 41). Also, like other RNA granules, the ORB proteome is enriched in proteins with intrinsically disordered regions and prion-like domains (Fig. 4A). ORB proteins form networks of physical and genetic interactions which are denser than would occur at random; 4.7 physical interactions per protein (p = 3.83 × 10^−9^) and 6.1 genetic interactions (p = 1.76 × 10^−6^) (Fig. 4B). Finally, ORBs have several members of the Ccr4-Not complex, which contributes to most aspects of RNA metabolism (Fig. S5, Table S1) (42). Thus, the validated proteome has concerted biochemical activities that would be expected in an organelle for RNA metabolism.

**Figure 4.**
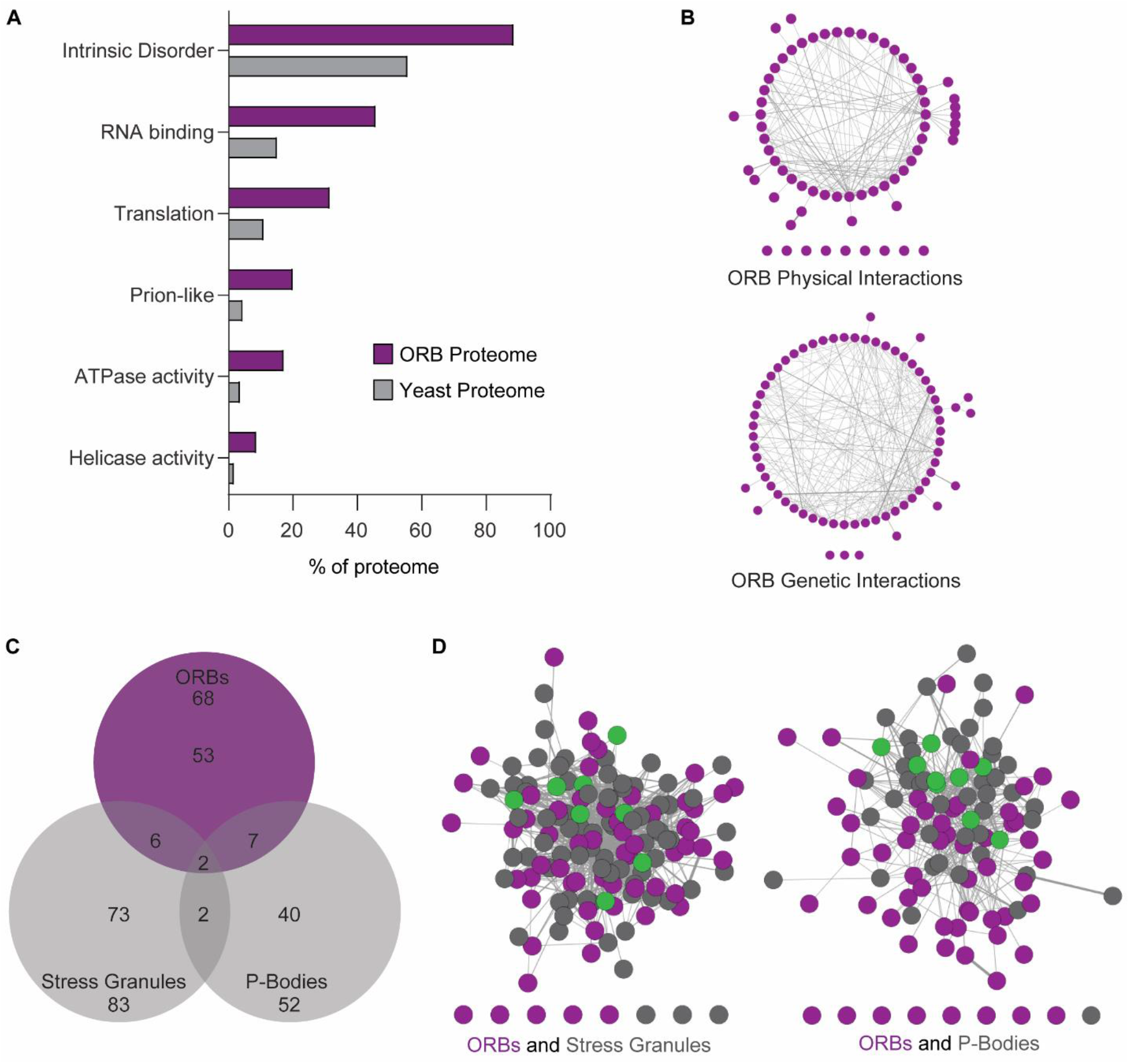
Features of the ORB proteome. (A) Enriched categories identified in the ORB proteome are presented as their percentage of the ORB proteome and, to show enrichment, the yeast proteome. (B) Known physical and genetic interaction networks within the ORB proteome. (C) Venn diagram showing limited overlap between the ORB proteome and the P-body and stress granule proteomes (Jain. et. al., 2016). (D) Known physical interactions between proteins in the ORB proteome and the proteome of either stress granules (left) or P-bodies (right). Shared proteins are shown in green.

A role of ORBs in the compartmentalization of TQC is supported further by the proteome. In addition to the TQC factors seen to localize to ORBs (Dom34, Hbs1, Hel2, Rqc2, (Fig.3A)), we identified four additional TQC proteins: Not4, Cdc48, Cue3, and Rtt101 (35, 43). Finally, a total of six ribosomal proteins of both subunits were identified, as expected for a TQC compartment (see above, Table S1).

### ORBs are Distinct from P-bodies and Stress Granules but Share Some Proteins with Each

This proteome allowed us to compare and contrast ORBs with P-bodies and stress granules, whose proteomes have been reported (18, 19). Comparisons revealed considerable non-overlap; most proteins in the ORB proteome are in neither P-bodies nor stress granules (Fig. 4C). During validation of the proteome, ORBs did not IF-stain positive for 2 additional P-body proteins (Ccr4, Xrn1) and 6 additional stress granule proteins (Hrp1, Pab1, Rbg1, Rio2, Rpo21, Rvb1) (Table S1). Therefore, ORBs are distinct from stress granules and P-bodies.

Nevertheless, ORBs appear to be functionally related to P-bodies and stress granules, based on partial overlaps of their proteomes (Fig. 4C). The ORB proteome has 8 stress granule proteins and 9 P-body proteins (Fig. 4C, Table S1). Two proteins are common to all three proteomes. Finally, analyses of the ORB proteome combined with either the stress granule proteome or the P-body proteome revealed dense interaction networks which involve many inter-proteome interactions (Fig. 4D). Therefore, OBS appear to be functionally related to stress granules and P-bodies, consistent with our evidence for their roles in mRNA metabolism and translation.

## DISCUSSION

### ORBs are Cytoplasmic RNA Granules

ORBs are RNA granules based on the following: they are cytoplasmic bodies that contain RNA; they are sensitive to 1-6,-hexanediol *in vivo*; they are resistant to the non-ionic detergent Triton X-100 *ex vivo* and, therefore, they are membaneless (Fig. 1) (26, 27). ORBs share functional classes of proteins with P-bodies and stress granules, e.g. proteins of RNA metabolism and the remodeling of RNP complexes, RNAs and proteins (Fig. 4) (18, 19). ORBs are enriched in proteins with intrinsically disordered regions and prion-like domains, both hallmarks of RNA granules. Finally, ORB proteins form a dense network of physical interactions, consistent with their being an organelle. Thus, ORBs are in a rapidly growing class of cytoplasmic RNA granules that compartmentalize processes related to translation (39, 44, 45).

### ORBs Compartmentalize Steps in TQC

We show that ORBs compartmentalize steps in TQC with four lines of evidence. First, active translation is required by both TQC and for the normal number of ORBs per cell (Fig. 2). Second, two distinct NGD substrates localize to ORBs; oxidized RNA and arrested RNCs (Figs. 1, 2). Third, NGD and RQC factors, as well as both ribosome subunits, localize to ORBs (Figs. 2, 3, 4). These factors function in the recognition of the arrested RNC (Hel2 and Cue3), the separation of the stalled ribosome subunits (Dom34, Hbs1, Rli1), the extraction of the nascent polypeptide from the 60S subunit (by CAT-tailing) and its ubiquitination (Rqc2, Cdc48, Not4) (34, 35). Fourth, mutants deficient for TQC factors have altered ORB numbers per cell or size (Fig. 3). Thus, the oxidized RNAs in ORBs could be mRNA substrates of TQC. Notably, ORBs appear to lack proteins involved in the endonucleolytic cleavage of the aberrant mRNA (Cue2) or the exonucleolytic degradation of the products (Xrn1, exosome subunits) (3, 46) (Table S1). The presence of Rtt101 and ribosome subunits in ORBs also suggests that aberrant rRNAs undergo quality control in ORBs (43). Additional work is required to address this possibility.

The localization of TQC to ORBs could have any of the known functions of intracellular compartmentalization, i.e., elevated local concentrations of substrates and intermediates to favor forward reactions and sequestration of aberrant molecules to prevent them from undergoing deleterious side-reactions. For example, truncated nascent polypeptides translated from aberrant mRNAs are prone to aggregation (1, 47–49). Sustained translation initiation on an oxidized mRNA could enhance the generation of aggregation-prone polypeptides. The formation of ORBs with Dom34 and Hbs1 *in vivo* likely reflects an elevated requirement for compartmentalized oxidized RNA quality control during H_2_O_2_-induced oxidative stress (Fig. 3 C-E).

The ORB proteome reveals candidate TQC proteins. For example, it has four ribosome nuclear export factors, which prevent association of unassembled ribosomal subunits until they are competent for translation (50). Subunit separation is required in TQC when the mRNA and nascent polypeptide are extracted from the disassembled ribosome (51). The ORB proteome includes four nuclear export factors; 2 for each ribosome subunit (40S, Nob1 and Tsr1; 60S, Lsg1 and Nmd3) (Table S1). These could be acting like Rqc2 by separating ribosome subunits during RQC (11, 52, 53). Therefore, our results open avenues to dissect TQC.

### The Position of ORBs Among the Cytoplasmic RNA Granules in Yeast

Our results support a role of ORBs in the TQC of non-translatable defective mRNAs with arrested ribosomes (Fig. 2). By contrast, translation granules are sites of active translation, Not1-containing assemblysomes handle temporarily paused ribosomes and P-bodies and stress granules handle translationally repressed mRNAs (39, 45, 54). Distinctions between ORBs and both P-bodies and stress granules are revealed by our results. This was particularly important because it has been reported that P-bodies are a location of NGD, and stress granules contain TQC substrates in a non-canonical stress-activated RQC pathway (16, 17). Only one TQC factor was identified in the proteome of stress granules (Rqc2) and none were identified in the proteome of P-bodies (18, 19). We show that ORBs lack at least 3 P-body proteins and 7 stress granule proteins (Table S1). These include canonical proteins of P-bodies (Dcp2, Xrn1) and stress granules (Pab1, Pub1). In addition, large majorities of the proteins in the ORB proteome are absent from the proteomes of both P-bodies and stress granules (Fig. 4). Ribosomal subunit localization also differs between these RNA granule types; P-bodies lack both subunits, stress granules have only the small subunit, and ORBs have both subunits (Fig. 2, Table S1) (18, 19). Puromycin stabilizes P-bodies and stress granules but decreased the number of ORBs per cell (Fig. 2) (28, 29). ORBs probably are not translation bodies because cycloheximide has opposite effects on their respective numbers per cell (Fig. 2) (39). P-bodies and stress granules have roles in the dynamic handling of mRNAs cycling to and from the translated pool on polysomes. These roles were revealed, in part, by the opposite effects on their formation when the initiation and elongation phases of translation are inhibited by puromycin and cycloheximide, respectively. By contrast, our finding that both cycloheximide and puromycin reduce ORB number suggests that defective mRNAs do not exit ORBs and return to polysomes, consistent with their degradation by TQC. Thus, ORBs are in a rapidly growing class of cytoplasmic RNA granules that compartmentalize processes related to translation (39, 44, 45).

Despite these differences, relationships between ORBs and both stress granules and P-bodies were revealed by partial overlaps of their proteomes. ORB localization was found for 9 P-body proteins and 8 stress granule proteins (Table S1, Fig. 4). Two proteins are common to P-bodies, stress granules, and ORBs: Dhh1 and Scd6. Analysis of the ORB proteome combined with the proteome of either P-bodies or stress granules revealed dense networks of physical interactions, supporting further functional relationships between ORBs and these RNA granules (Fig. 4). In addition, ORB and P-body proteins partitioned to separate sectors within the combined physical interactome (Fig. 4D). This segregation has been seen for P-body and stress granule proteins (18,19). By contrast, the ORB and stress granule proteomes are more interconnected, suggesting a close functional integration.

## METHODS

### Plasmids, Yeast Strains, and Growth Conditions

Yeast plasmids were constructed using standard molecular biology procedures starting from p415ADH1 (56) and were described previously (Navickas et al., 2020). pLB138 expresses a truncated mRNA (Rz) from a 2HA-tagged-*URA3*-RZ (Ribozyme) construct. pLB126 expresses a full length 2HA-*URA3* mRNA (FL). The wild-type strain was BY4741 (*MATa his3Δ1 leu2Δ0 met15Δ0 ura3Δ0*). All single mutant strains were obtained from the yeast knockout collection in the BY4741 background (57). All strains with GFP-tagged proteins were obtained from the GFP clone collection in the BY4741 background (58). The *dom34*Δ-*rqc2*Δ double mutant was generated by one-step gene replacement using PCR fragment of the NatMX6 cassette amplified from plasmid pFA6a-natMX6 (59).

Strains were grown at 30°C in YPD to mid-log phase (OD600 of 0.5-0.8). Where indicated, cells were exposed to 100 μg/ml cycloheximide, 100 μg/ml puromycin, or 5% 1,6-hexanediol with 10 μg/ml digitonin for 30 min. For stress conditions, cells were exposed for 30 min to either: YPD without glucose, 0.5% sodium azide, or 5 mM hydrogen peroxide. To induce autophagy, cells in mid-log phase were transferred to nitrogen starvation minimal medium (SD-N) for 2 hrs. Where indicated, cells were treated with 3.7% formaldehyde for 30 minutes.

### Western Blotting

Yeast cells from mid-log phase cultures were incubated on ice for 10 min in lysis buffer (0.2 M NaOH, 0.2% β-mercaptoethanol), whereupon 5% trichloroacetic acid was added followed by a 10-min incubation on ice. Precipitated proteins were pelleted by centrifugation at 12,000 × g for 5 min and resuspended in 35 μL of dissociation buffer (4% SDS, 0.1 M Tris-HCl, pH 6.8, 4 mM EDTA, 20% glycerol, 2% β-mercaptoethanol and 0.02% bromophenol blue). Tris-HCl, pH 6.8 was then added to a final concentration of 0.3 M and samples were incubated at 37°C for 10 min. Total protein extracts were subjected to SDS-PAGE and immunoblot analysis. Membranes were reacted with 1:2000 mouse αHA primary antibody (Covance). Secondary staining used 1:10,000 goat anti-mouse-IgG antibody (KPL).

### Indirect IF

Yeast cells from mid-log phase cultures were fixed in 3.7% formaldehyde for 15 min at room temperature, washed in phosphate buffer (0.1 M potassium phosphate, pH 6.5), and incubated with 10 mg/ml lyticase (in phosphate buffer containing 1.2 M sorbitol) for 30 min at 30°C. Spheroplasts were adhered to 0.1% poly-L-lysine-coated slides for 10 min. Slides were then submerged in ice-cold methanol for 5 min followed by room-temperature acetone for 30 seconds. Cells were blocked in 2% BSA, 1X PBS, 0.1% Tween-20 for 10 min and incubated with primary antibody (1:500 mouse α8-oxoG, QED Bioscience; 1:5000 rabbit αGFP, ThermoFisher, 1:5000 mouse αHA, Covance) overnight at 4°C in a humidity chamber. Secondary antibodies were Alexa Fluor 647 donkey αgoat IgG, Alexa Fluor 568 goat αmouse IgG, and Alexa Fluor 488 goat αrabbit IgG, each diluted 1:300 (ThermoFisher). The presence of RNA in ORBs was detected by staining with a 1:500 dilution of SYTO RNASelect™ (ThermoFisher) for 10 min prior to mounting. Where indicated, fixed and permeabilized cells were treated for 1h at room temperature with 10 μg/ml RNase A (Fermentas) prior to blocking. The specificity of the α8-oxoG antibody was confirmed by incubating with its antigen 8-hydroxy-2′-deoxyguanosine (10 μg/μg of antibody) for 2 h before IF-staining; the αHA antibody is specific because the signal was not seen in a non-transformed strain or when the primary αHA antibody is omitted in IF of a pLB126 transformant, and it was significantly lower in the *dom34*Δ mutant; The GFP antibody is specific because no signal was seen in a non-transformed strain (Fig. S6).

### Preparation of the ORB-Enriched Fraction

ORBs were prepared from BY4741 cells or the indicated strains from the GFP fusion library (58), as described previously for stress granule cores (19). Briefly, cell pellets from 50 mL cultures grown to mid-log phase were resuspended in 500 μL lysis buffer (50 mM Tris-HCL (pH 7.5), 50 mM NaCl, 5 mM MgCl_2_, 50 μg/mL heparin, 0.5 mM DTT, 0.5% Nonidet P-40, fungal protease inhibitor (BioShop), 1:5000 Antifoam B (Sigma) and lysed by vortexing with acid-washed glass beads for 2 min followed by 2 min on ice for three cycles. The initial lysate was cleared of unbroken material by centrifugation at 800 × g for 2 min at 4°C, and then centrifuged at 17,200 × g for 10 min at 4°C. This ORB-enriched pellet fraction was then resuspended in a final volume of 500 μL lysis buffer on ice. To test RNase A and Triton X-100 sensitivity of ORBs *ex vivo*, the final pellet was resuspended in lysis buffer containing either 10 ug/mL RNase A without heparin or 2% Triton X-100 and incubated for 30 min at room temperature on a nutator before a second centrifugation at 17,200 × g for 10 min at 4°C and resuspension in fresh lysis buffer. ORBs were detected by adhering the ORB-enriched fraction to poly-L-lysine-coated slides and probing for 8-oxoG as described above.

### Microscopy and Image Analysis

Images were acquired using a Leica DMI 6000 epifluorescence microscope (Leica Microsystems) with a 63X/1.4NA objective, a Hamamatsu OrcaR2 camera, and Volocity acquisition software (Perkin-Elmer). Z-stacks were taken by series capture at a thickness of 0.2 μm per section. Stacks were deconvoluted with AutoQuant X3 (Media Cybernetics Inc.). To quantify ORBs *in situ* we used a custom-written macro in ImageJ. Briefly, each cell was identified in a central z-slice, and its average 8-oxoG signal intensity determined. ORBs were defined as foci between 9 and 198 pixels^2^ with a fluorescence intensity 1.5-fold higher than the cell average. The number of ORBs per cell, their size, and fluorescence intensity were then quantified by the macro. To quantify ORBs *ex vivo* we used the 3D objects counter plugin in ImageJ. ORB colocalization of stalled RNCs or GFP-tagged proteins *in situ* was determined using a custom-written ImageJ macro. Briefly, foci in each channel were determined as described above, and two foci from different channels were considered colocalized if at least 25% of the area of each foci in each channel overlapped. The number and size of GFP foci in H_2_O_2_-and formaldehyde-treated live cells were measured with FIJI using the multi-point and line-selection tools, respectively (60). For a candidate protein to be considered a validated ORB protein, three criteria were required: visual inspection of colocalization must be positive; at least 10% of identified ORBs must colocalize with a GFP foci; at least 18% of cells must have at least one colocalized foci. A minimum of three images were quantified for each biological replicate. One replicate from WT-FL in Fig 2F was considered an outlier because it was above 2 SEM and was not considered.

### Purification of ORBs

ORBs were purified as described previously for stress granule cores (61). Protein A Dynabeads (Invitrogen) were added to ORB-enriched fractions from either untreated or cycloheximide-treated cultures for 1 hr at room temperature to pre-clear the fraction of non-specific interactions. Cleared lysates were then incubated with α8-oxoG (QED Bioscience) at 4°C for 1 hr to capture ORBs, followed by a 1 hr incubation with Dynabeads to capture the ORB-antibody complexes. Purified ORBs were resuspended and boiled in SDS-loading buffer for 5 min, then loaded onto a 5% polyacrylamide gel for in-gel trypsin digestion.

### Mass Spectrometry

Purified ORB proteins were concentrated in a stacking gel using SDS-PAGE stained with Coomassie Brilliant Blue R-250 (Biorad). Proteins in the excised gel band were subjected to in-gel digestion as follows. Gel pieces were incubated for 30 min at room temperature in 50 mM NH_4_HCO_3_ (Sigma) + 10 mM dithiothreitol to reduce the proteins and then for 30 min at room temperature with 50 mM NH4HCO3 + 50 mM Iodoacetamide (SIGMA) in the dark to alkylate them. Gel dehydration was done with a series of acetonitrile (ACN, BDH) washes. Gel pieces were rehydrated in trypsin digestion solution containing 25 mM NH4HCO3 and 10 ng/μL of trypsin (Sigma) followed by incubation overnight at 30°C. Tryptic peptides were extracted three times for 15 min at room temperature with extraction solution (60% acetonitrile + 0.5% formic acid (FA, Fisher), four volumes of the digestion solution). Peptides were dried using a Speedvac at 43°C and stored at −20°C until MS analysis. Liquid chromatography-tandem MS (LC-MS/MS) analyses were performed on a Thermo EASY nLC II LC system coupled to a Thermo LTQ Orbitrap Velos mass spectrometer equipped with a nanospray ion source. Tryptic peptides were resuspended in solubilization solution containing 97% of water, 2% of ACN and 1% of FA to give a peptide concentration of 100ng/μL. 2 μL of each sample were injected into a 10 cm × 75 μm column that was in-house packed with Michrom Magic C18 stationary phase (5 μm particle diameter and 300 Å pore size). Peptides were eluted using a 90-min gradient at a flow rate of 400 nL/min with mobile phase A (96.9% water, 3% ACN and 0.1% FA) and B (97% ACN, 2.9% water and 0.1% FA). The gradient started at 2% of B, linear gradients of B were achieved to 8% at 16 min, 16% at 53 min, 24% at 69 min, 32% at 74 min, 54% at 81 min, 87% at 84 min followed by an isocratic step at 87% for 3 min and at 2% for 3 min. A full MS spectrum (m/z 400-1400) was acquired in the Orbitrap at a resolution of 60,000, then the ten most abundant multiple charged ions were selected for MS/MS sequencing in linear trap with the option of dynamic exclusion. Peptide fragmentation was performed using collision induced dissociation at normalized collision energy of 35% with activation time of 10 ms. Spectra were internally calibrated using polycyclodimethylsiloxane (*m/z* 445.12003 Da) as a lock mass.

### Mass Spectrometric Data Processing

The MS data were processed using Thermo Proteome Discoverer software (v2.2) with the SEQUEST search engine. The enzyme for database search was chosen as trypsin (full) and maximum missed cleavage sites were set at 2. Mass tolerances of the precursor ion and fragment ion were set at 10 ppm and 0.6 Da, respectively. Static modification on cysteine (carbamidomethyl, +57.021 Da) and dynamic modifications on methionine (oxidation, +15.995 Da) and N-terminus (acetyl, +42.011 Da) were allowed. The initial list contained 1048 proteins identified with high confidence (false discovery rate <1%) and from combining three biological replicates. Of this list we retained 822 proteins according to criteria stated in the results section.

### Bioinformatic Analysis of ORB proteome

The ORB proteome was classified according to GO Molecular Function and GO Biological Process information from the SGD database (yeastgenome.org). RNA binding activity was determined using GO analysis and published lists of yeast RNA binding proteins (40, 41). Networks of physical and genetic interactions were generated and visualized using Cytoscape (version 3.7.2) with the GeneMania plugin (version 3.5.2) (62, 63). Proteins with intrinsically disordered regions or prion-like domains were predicted using the SLIDER and PLAAC tools, respectively (64, 65).

## END NOTES

## Acknowledgments

For technical support, materials, infrastructure and advice, we thank Dr. Heng Jiang and the Centre for Biological Applications of Mass Spectrometry, Dr. Christopher Law and the Centre for Microscopy and Cell Imaging, Dr. Christopher Brett (Concordia University) and the members of his laboratory, and the Centre for Structural and Functional Genomics. This work was supported by Discovery Grant (217566) from the Natural Sciences and Engineering Research Council of Canada to WZ.

## Author Contributions

J.S.D., L.B., and W.Z. designed the study, analyzed the data. J.S.D. and W.Z. wrote the initial draft; J.S.D. performed experiments and collected data; C.P. and L.B. provided resources; J.S.D., C.P., L.B., and W.Z revised the manuscript.

## Supplementary Information

Supplemental Figures: PDF containing 6 supplementary figures

Table S1: .xlsx file containing the list of candidate and validated ORB proteins

## Code Availability

ImageJ macros used to quantify images are accessible at https://github.com/Zergeslab/

## SUPPLEMENTARY FIGURES

**Figure S1.**
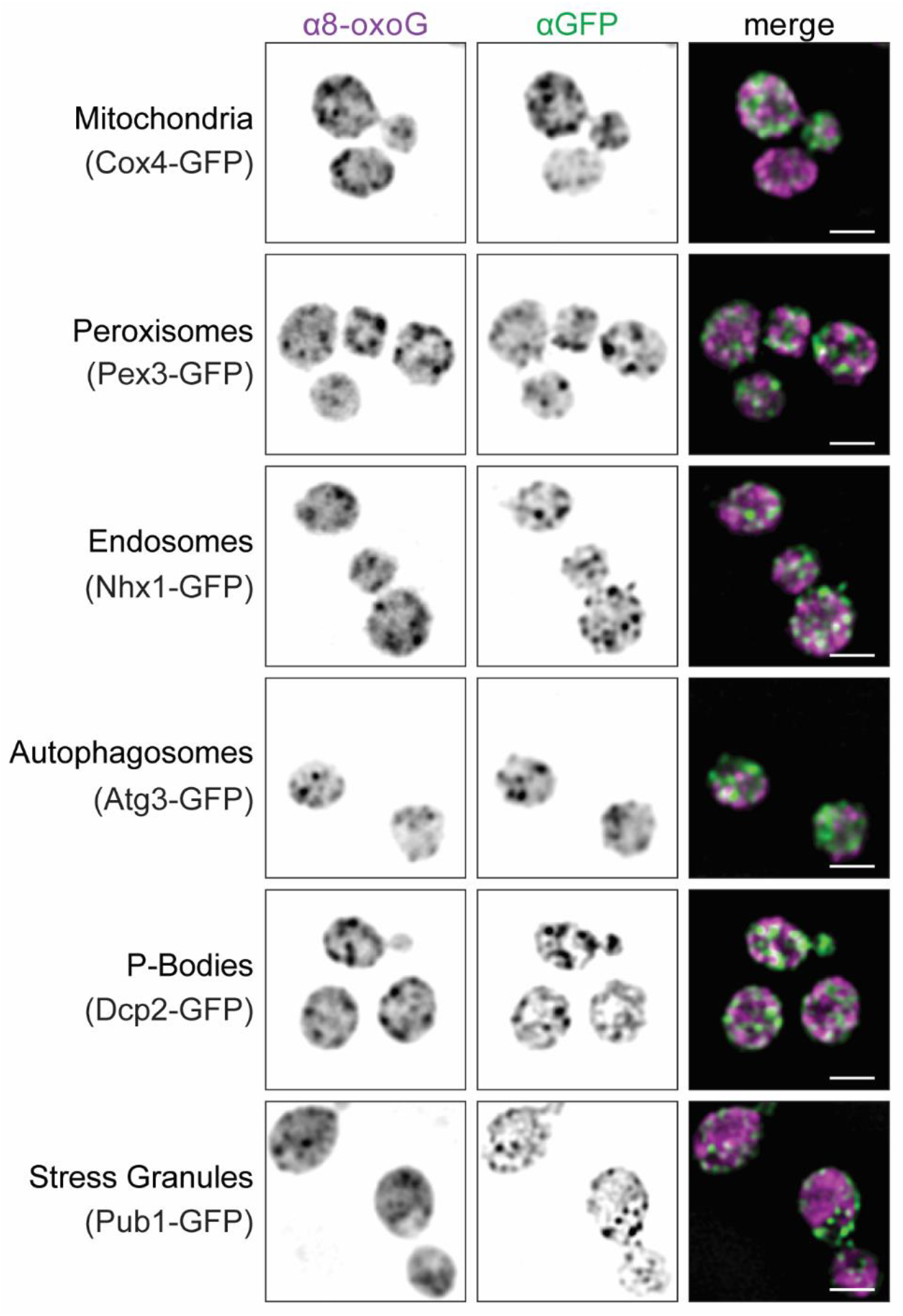
ORBs are none of the known RNA-containing organelles. Co-IF staining of ORBs (α8-oxoG) and the indicated GFP-tagged marker proteins for membranous organelles and RNA granules *in situ*. Scale bars indicate 2.0 μm.

**Figure S2.**
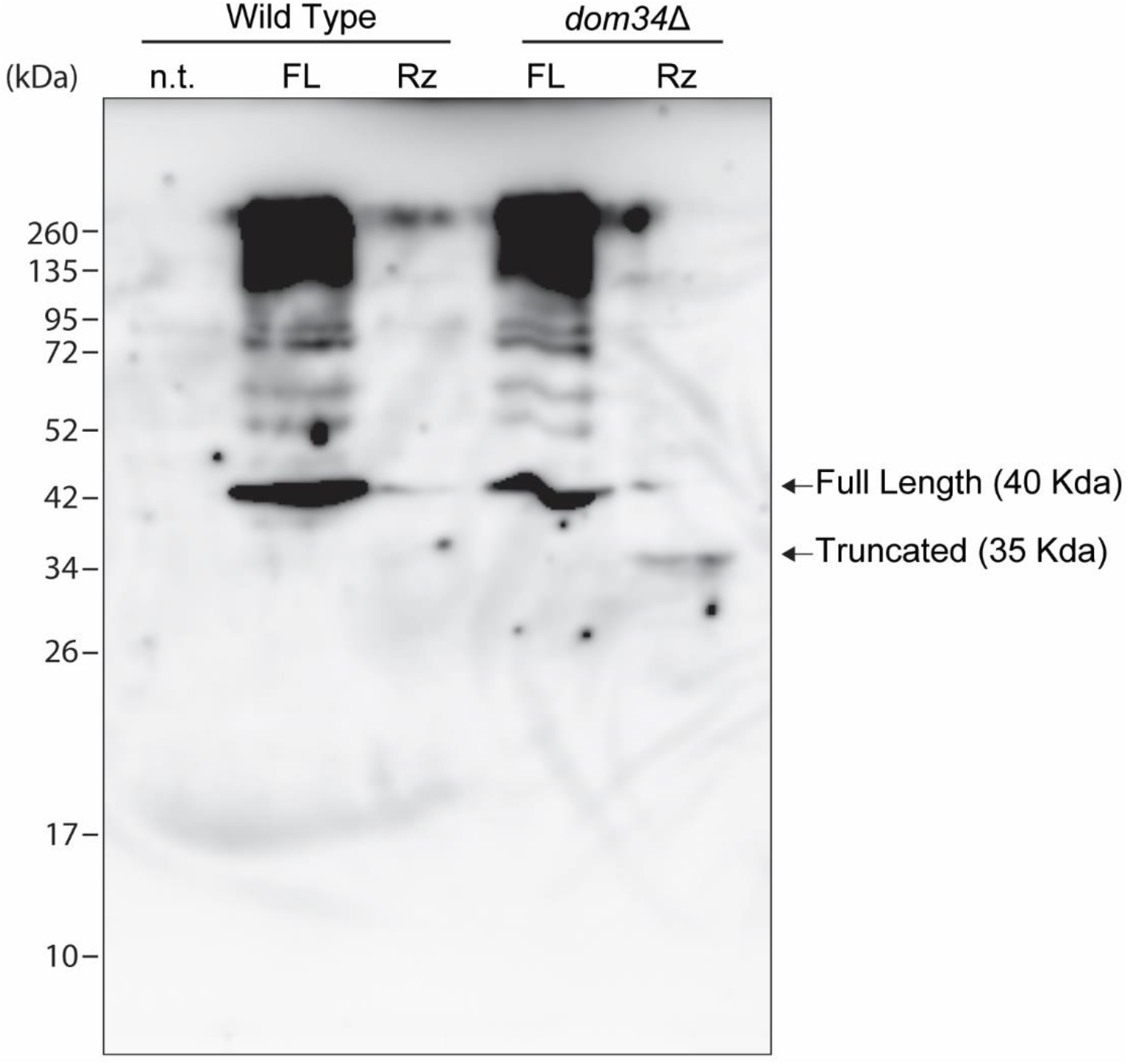
The truncated HA-Ura3 nascent polypeptide accumulates in *dom34*Δ background. Immunoblot analysis of HA-Ura3 in total protein from either WT or *dom34*Δ, transformed with the expression construct for the HA-*URA3* mRNA, either with the self-cleaving hammerhead ribozyme (Rz) or without it (FL). n.t.=non-transformed WT.

**Figure S3.**
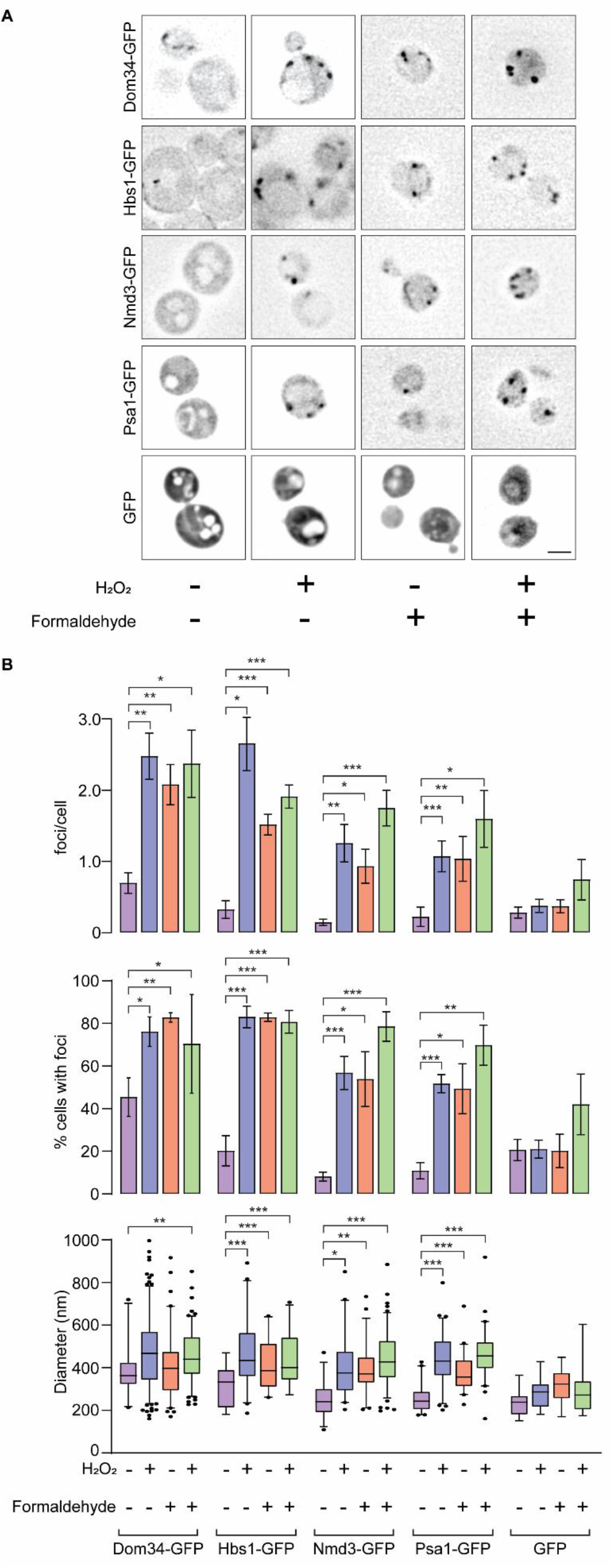
ORB proteins form foci in response to H_2_O_2_ and formaldehyde. (A) Dom34-GFP, Hbs1-GFP, Nmd3-GFP, Psa1-GFP, and GFP fluorescence *in vivo*, without or with exposure to 5.0 mM H_2_O_2_, 3.7% formaldehyde, or both. (B) Graph of foci per cell (top), percentage of cells with foci (middle), and foci diameter (bottom) for the proteins indicated in (A), without or with exposure to H_2_O_2_. Bar heights indicate the average (n=4). Error bars = ± 1 SEM. Statistical significance was determined by unpaired two-sample t-tests. Box and whisker plots indicate the interquartile range, median values, and the 5^th^ and 95^th^ percentiles from all foci measured across the four replicates. Scale bars = 2.0 μm. N values refer to the number of independent biological replicate experiments. * = p ≤ 0.05, ** = p ≤ 0.01, *** = p ≤ 0.001.

**Figure S4.**
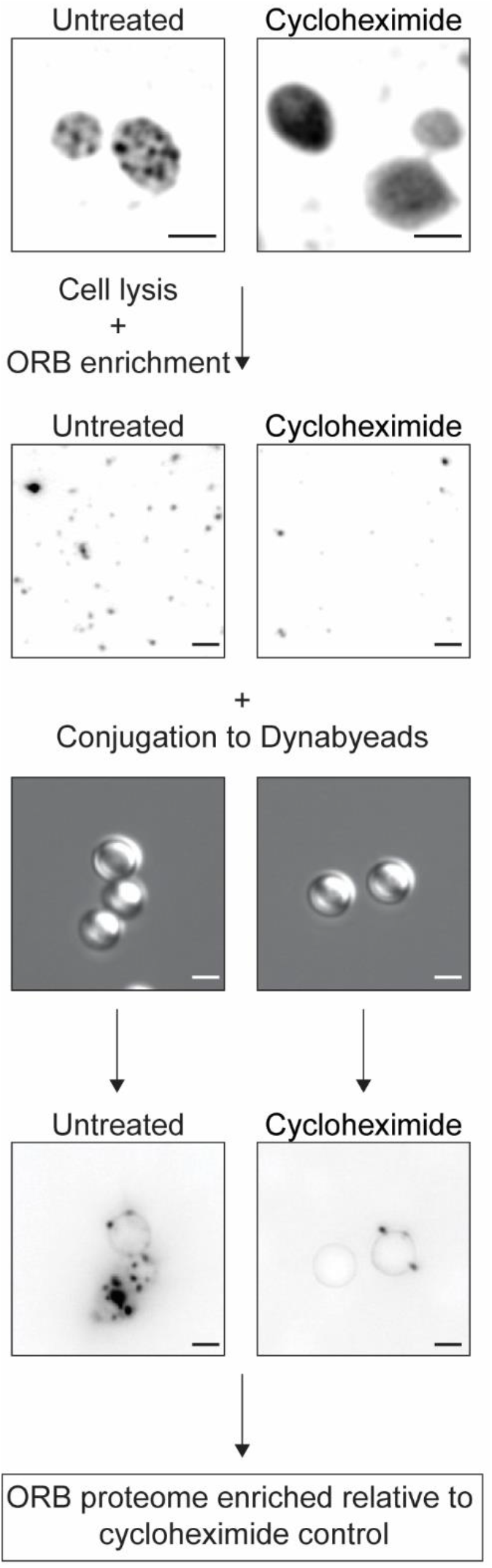
ORB enrichment strategy. The flow chart illustrates cell lysis and immunoprecipitation of 8-oxoG from cell pellets of untreated or cycloheximide-treated cells.

**Figure S5.**
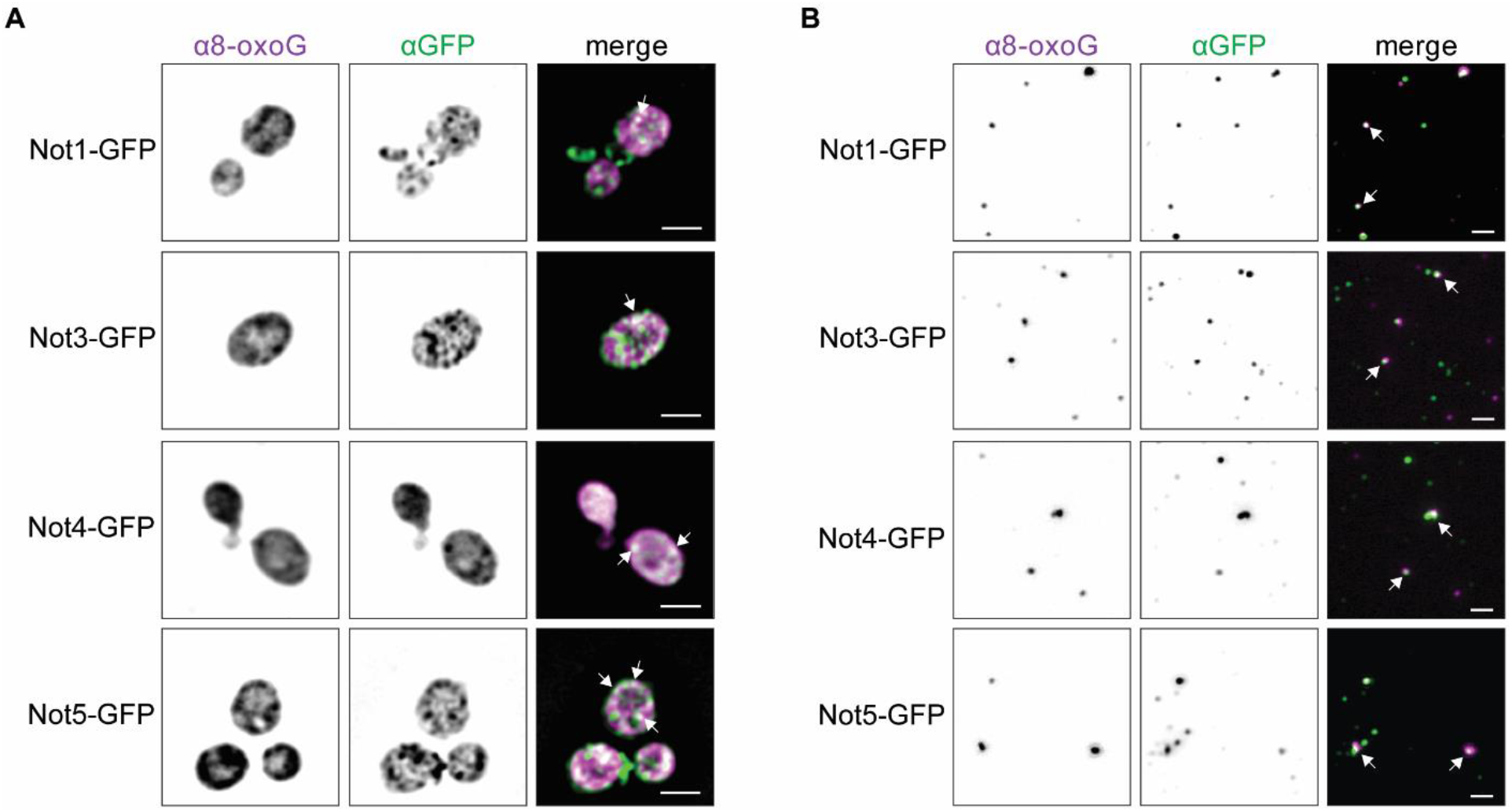
Components of the Ccr4-Not complex localize to ORBs. (A) *in situ* co-IF staining of ORBs (α8-oxoG) and the indicated GFP-tagged proteins *in situ*. (B) *ex vivo* co-IF staining of ORBs (α8-oxoG) and indicated GFP-tagged proteins. Arrows indicate foci of co-localized signals. Scale bars = 2.0 μm.

**Figure S6.**
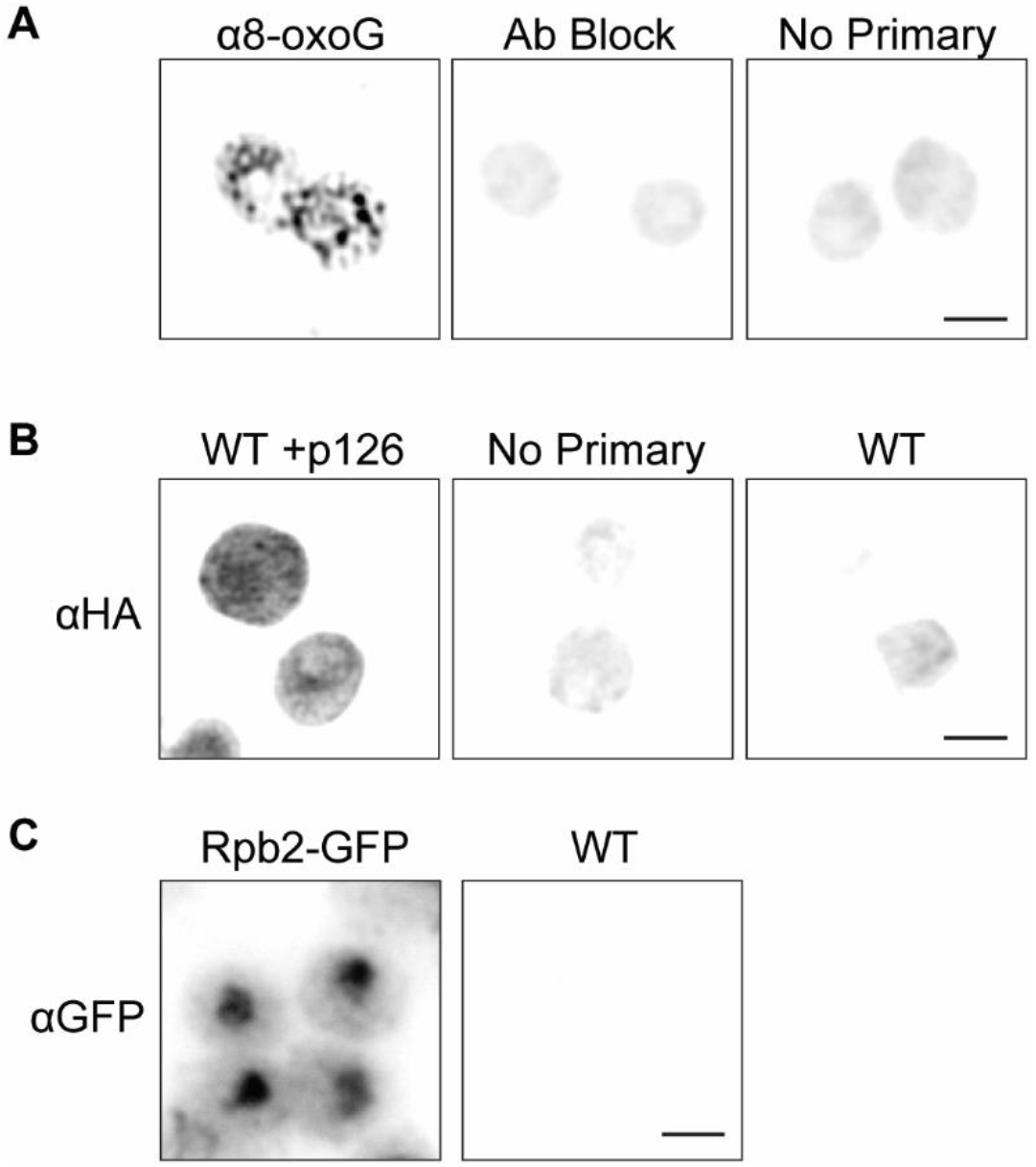
Antibody staining controls. (A) ORBs are not visible when the primary antibody against 8-oxoG was excluded (No Primary) or following blocking of the antibody with 8-oxoG (Ab Block). (B) No primary antibody control (No Primary) and staining of untransformed wild type cells (WT), demonstrating that the HA signal seen *in situ* is specific. (C) IF staining of GFP in untransformed cells (WT) or cells transformed with Rpb2-GFP, demonstrating that the GFP signal seen *in situ* is specific. Scale bars = 2.0 μm.

